# *BundleAGE*: Predicting White Matter Age using Along-Tract Microstructural Profiles from Diffusion MRI

**DOI:** 10.1101/2024.08.16.608347

**Authors:** Yixue Feng, Julio E. Villalón-Reina, Talia M. Nir, Bramsh Q. Chandio, Neda Jahanshad, Paul M. Thompson

## Abstract

Brain Age Gap Estimation (BrainAGE) is an estimate of the gap between a person’s chronological age (CA) and a measure of their brain’s ‘biological age’ (BA). This metric is often used as a marker of accelerated aging, albeit with some caveats. Age prediction models trained on brain structural and functional MRI have been employed to derive BrainAGE biomarkers, for predicting the risk of neurodegeneration. While voxel-based and along-tract microstructural maps from diffusion MRI have been used to study brain aging, no studies have evaluated along-tract microstructure for computing BrainAGE. In this study, we train machine learning models to predict a person’s age using along-tract microstructural profiles from diffusion tensor imaging. We were able to demonstrate differential aging patterns across different white matter bundles and microstructural measures. The novel Bundle Age Gap Estimation (BundleAGE) biomarker shows potential in quantifying risk factors for neurodegenerative diseases and aging, while incorporating finer scale information throughout white matter bundles.

## I. Introduction

Brain Age Gap Estimation (BrainAGE) is widely used as a metric of accelerated aging, and to predict a person’s risk of developing neurodegenerative diseases. BrainAGE makes the distinction between a person’s chronological age and an estimate of their biological age. The difference between these measures can be useful in predicting the risk of dementia, and all-cause mortality [1]. BrainAGE is often estimated from voxel-based brain MRI data using multivariate machine learning models. Studies have shown the association of BrainAGE with cognitive decline and dementia severity [2]. Interpretability analysis reveals spatial patterns that correspond to the profile of atrophy associated with healthy aging [3].

Diffusion MRI (dMRI) [4] is sensitive to alterations in white matter (WM) microstructure, which, in turn, is affected in neurodegenerative diseases such as Alzheimer’s disease. WM microstructure is typically studied using scalar metrics calculated at the voxel- or region of interest (ROI) level, such as those derived from diffusion tensor imaging (DTI) [5]. However, tractography can be used to reconstruct WM tracts as 3D geometric models of their trajectories, and these provide more detailed representations of their underlying structure. Tractometry is an approach that integrates tissue microstructural metrics with tractography reconstruction to map WM abnormalities along the length of the tract [6]–[ Using 3D convolutional neural networks (CNN), scalar maps from diffusion tensor imaging (DTI) have been employed for age prediction in diverse cohorts [9]–[11] However, few studies have evaluated the predictive performance of along-tract profiles for brain aging. Kruper *et al*. trained a 1D CNN model for glaucoma classification from along-tract microstructural profiles, and found that the visual pathways (optic radiation tracts) performed better than non-visual pathways [12]. However, no work has examined along-tract features for brain age analysis using tractometry approaches.

In this study, we propose the BundleAGE framework for age prediction using along-tract microstructural profiles of 4 widely used DTI metrics for 31 major WM tracts in the brain. Along-tract profiles serves as an effective middle ground between the high-granularity of whole-brain voxel-wise analysis, and the coarse resolution of measures averaged over larger ROIs.

## II. Methods

### A. Data

We analyzed data from 568 healthy controls (HC) from the Lifespan Human Connectome Project Aging (HCP-A) study (239 male / 329 female; 36-100 years old; mean age: 57.45*±* 14.07 (SD) years) [13]. Accquisition parameters for the diffusion and T1-weighted structural images can be found in [14]. Preprocessing of dMRI of includes denoising using local principal component analysis [15] and Gibbs ringing correction [16], [17] implemented in DIPY [18], followed by the HCP pipeline [19]. Diffusion images registered to the MNI space were used to fit diffusion tensors at each voxel using the non-linear least-squares method to produce four scalar maps of fractional anisotropy (FA), mean (MD), radial (RD), and axial (AxD) diffusivity. Fiber orientations were reconstructed using Robust and Unbiased Model-BAsed Spherical Deconvolution (RUMBA-SD) [20] with Contextual Enhancement [21]. Whole-brain tractograms were generated using particular filtering tracking [22], with 27 seeds per voxel generated from the WM mask, a step size of 0.75 mm, angular threshold of 30^*°*^, and the continuous map stopping criterion. BundleSeg [23] was used to segment 31 WM tracts using the population-averaged HCP-842 atlas [24].

The Bundle Analytics (BUAN) tractometry pipeline [7] was employed to create bundle profiles of four microstructural metrics along 50 tract segments aligned across subjects for 31 tracts. Namely, scalar maps of these metrics were mapped to each point on each streamline and averaged for each segment, see **Figure 1**. A truncated mean was computed discarding the top and bottom 10% of values for each segment to reduce the effect of outliers. This resulted in 50 features per DTI metric per subject to be used for the age prediction task.

**Fig. 1.**
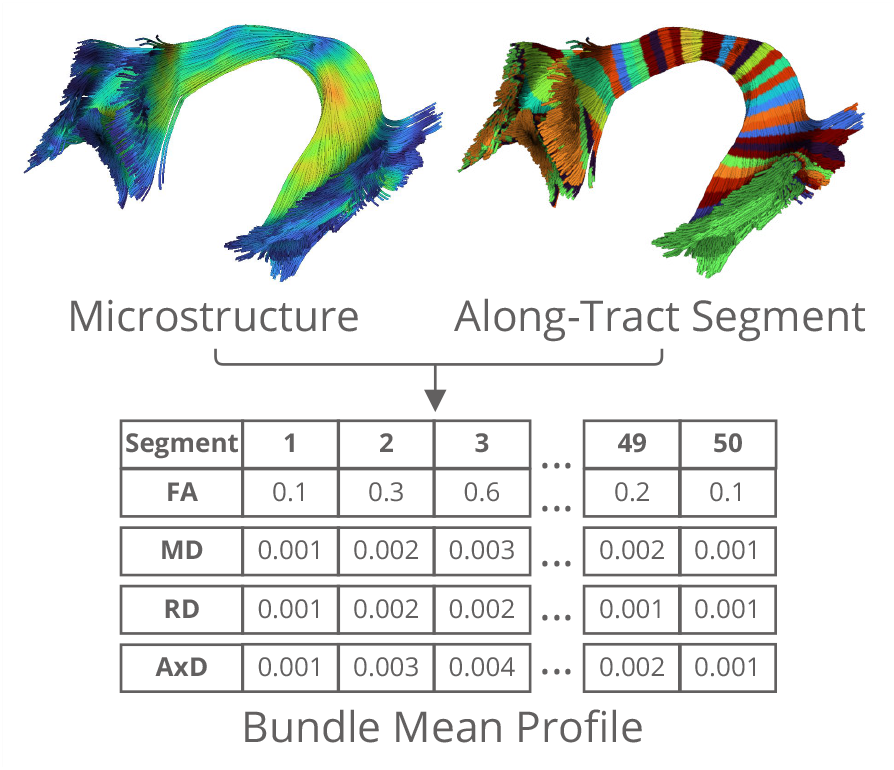
Scalar maps of DTI metrics are mapped to each point on a bundle. Along-tract segments are created using the atlas bundle as a reference, so that they are aligned across subjects. DTI metrics are averaged for all points that belong to each segment to create the bundle profile for downstream tasks.

### B. Age Prediction Model

We first created an 80/20 train/test split across all participants stratified by sex, resulting in 454 training and 114 test subjects. The 80% training split was then used to train a random forest (RF) regression model (implemented in scikit-learn v1.1.1) to predict the chronological age, measured in months. For each bundle and DTI metric, we performed a grid search over the following parameters using 5-fold cross-validation (CV) on the 80% training data: the maximum depth of the tree (5, 10, 15), the number of trees in the forest (100, 250, 500), the maximum number of features used for split (7, 20, 35, 50), and the minimum number for a leaf node (1, 3, 5). Training data for each CV fold was standardize to have zero mean and unit variance, and the same scaler parameters were applied to the remaining data. The coefficient of determination (*R*^2^) was used to evaluate each model during grid search, and the best performing model was then applied to the 20% held-out test set for evaluation. In addition to *R*^2^, we also calculated the mean absolute error (MAE) and Pearson correlation coefficient (*r*) between the model predicted biological age and chronological age.

In addition training the RF model on 50 along-tract features for each DTI metric, we concatenated features from mean profiles 4 DTI metrics to create 200 features per bundle, and trained an additional model using the same procedures described above.

## III. Results

Results of three evaluation metrics – *R*^2^, MAE, and *r* – for the age prediction model of bundle profiles, are shown in **Figure 2**. For all bundles and metrics, the predicted brain age values are significant correlated (*p*_FDR_ *<* 0.05) with the chronological age, after controlling for multiple test with FDR correction. Out of all bundles, models trained on inferior fronto-occipital fasciculus left (IFOF L), cingulum left (C L) and optic radiation right (OR R) achieved the best performance on three evaluation metrics. Models trained on the spino-thalamic tracts (STT R) in the brainstem had the worst performance, along with the middle longitudinal fasciculus right (MdLF R).

**Fig. 2.**
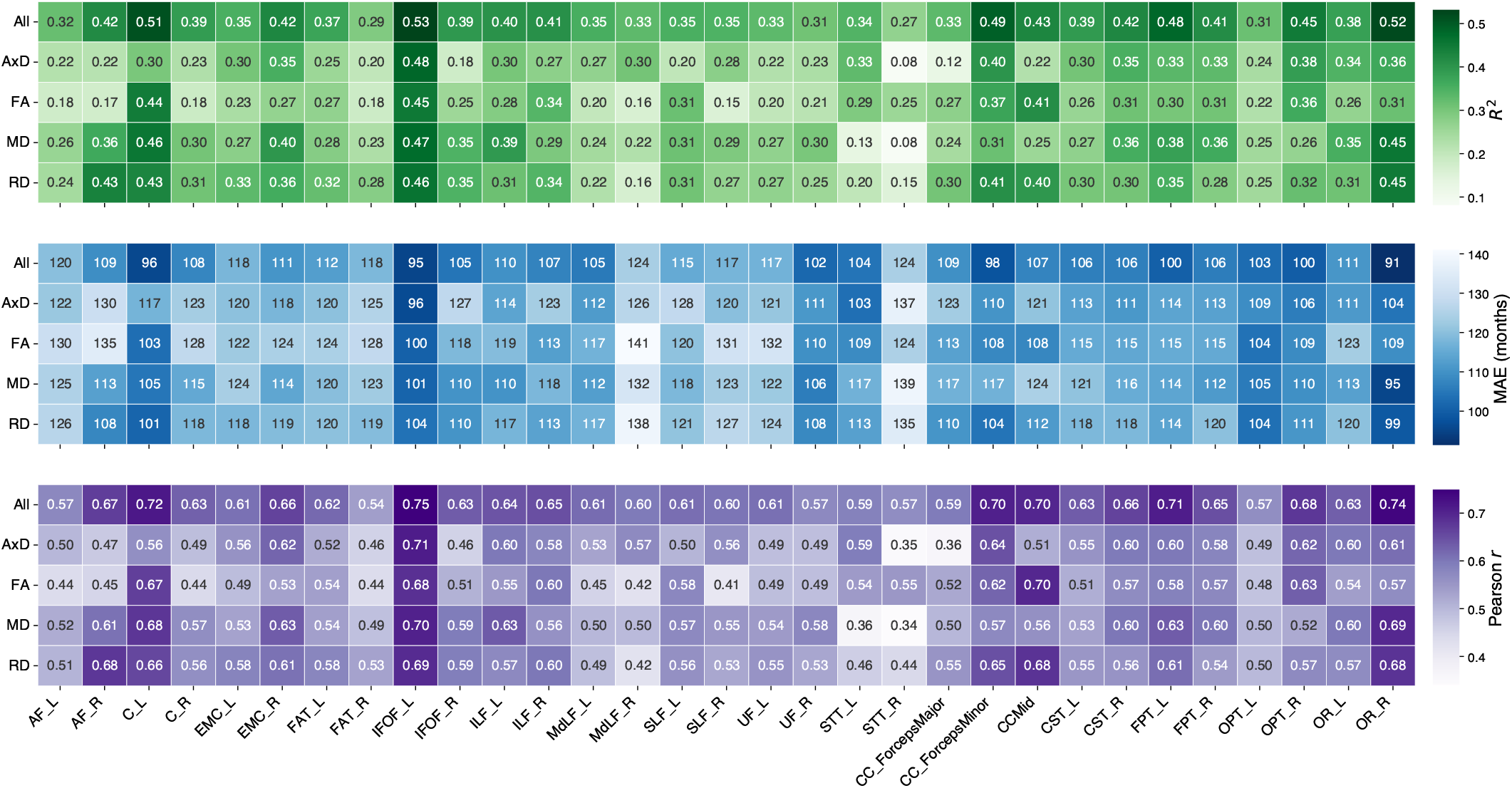
Coefficient of determination (R2), mean absolute error (MAE), and Pearson’s correlation coefficient (r) for each random forest regression model trained on the microstructural profile of 31 WM bundles, which include features from AxD, FA, MD, RD and all concatenated features.

Models trained on concatenated features outperformed single metric features except for RD of AF R, AxD of STT L. The DTI metrics used for age prediction reflect different characteristics of water diffusion, and when combined, can provide more comprehensive information for the model. For models trained on bundle profiles of single metric, MD and RD performed better than FA and AxD on average, consistent with prior literature examining the relationship between regional DTI measures with age [25], [26]. The best performing model out of all experiments is the along-tract concatenated profile of OR R, which achieved an MAE of 91 months (7.58 years) on the test set.

The random forest regression model produces an impurity-based feature importance score, in which a higher value corresponds to a more important feature (bundle segment in our experiments). We plot the along-tract feature importance scores for the three best performing bundles from models trained on the concatenated features in **Figure 3**. For these bundles, patterns of feature importance are similar across DTI measures, with AxD being the most dissimilar from the other measures. Salient features are concentrated in the frontal regions of the C L and IFOF L bundle, and are more widespread than salient features for OR R.

**Fig. 3.**
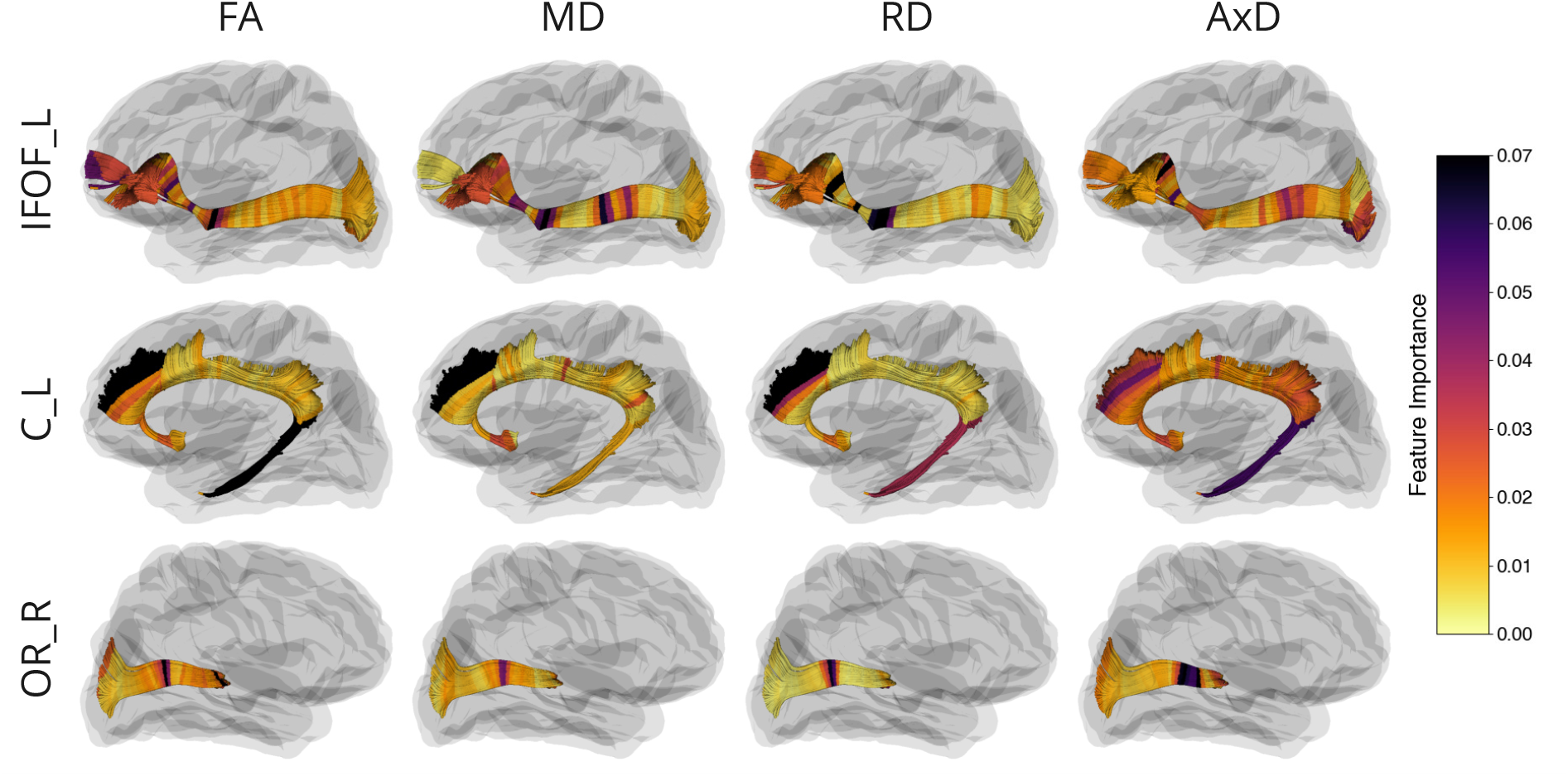
Along-tract feature importance for 4 DTI measures of the three best performing bundles from models trained on the concatenated features.

Due to the age-dependent bias of the brain age gap [27], we used correlation-constrained linear regression to calculated a corrected brain age [28]. Scatter plots of the chronological age versus the corrected biological age of the two best performing models trained on single metric features are shown in **Figure 4**.

**Fig. 4.**
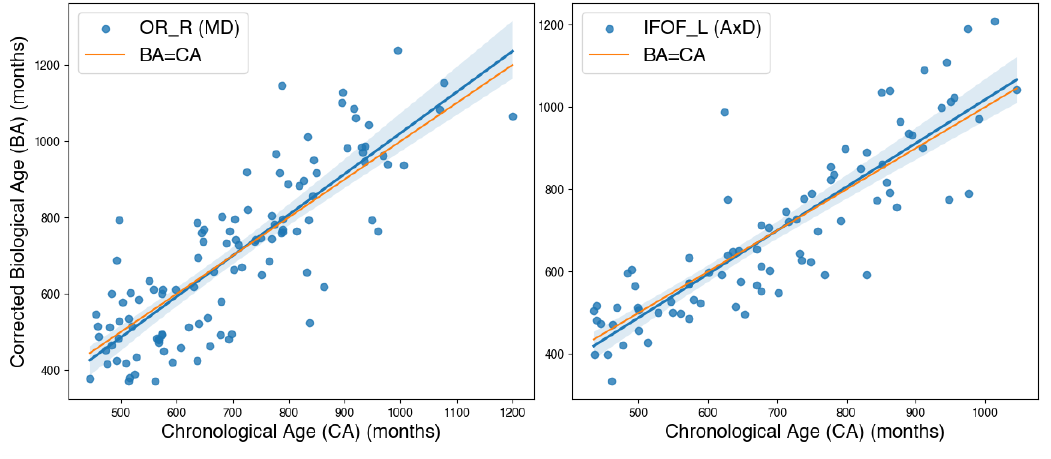
Comparison of chronological age (CA) and biological age (BA) for the two best performing models trained on single metric features.

## IV. Conclusions and Future Work

In this study, we propose bundle-specific brain age prediction using along-tract microstructural profiles from 31 major WM bundles. Machine learning models trained on these profiles achieve the best performance for the right optic radiation (OR R), left inferior fronto-occipital fasciculus (IFOF L), and left cingulum (C L), as well as the corpus callosum. Although models trained on concatenated features performed the best, profiles of MD and RD, which characterize diffusion in non-principal directions, outperform AxD and FA, consistent with prior literature. By retaining spatial variability of tissue microstructure along WM tracts that are often lost in ROI-based methods, while avoiding the computational challenges associated with 3D whole-brain models, our approach allows for a practical and nuanced approach for studying the role of WM in aging.

In our future work we will calculate and evaluate BundleAGE on additional clinical datasets, such as the Alzheimer’s Disease Neuroimaging Initiative (ADNI) [29], which includes participants with mild cognitive impairment and dementia. We will relate the BundleAGE biomarkers to neurocognitive assessments of dementia severity as well as measures of amyloid and tau. With a larger sample, we will also include additional microstructural measures from advanced diffusion models such as neurite orientation dispersion and density imaging (NODDI) [30] and diffusion kurtosis imaging (DKI) [31], and examine their effect on BundleAGE.

## Acknowledgment

This work was supported by the National Institutes of Health under the FiberNET project grant, RF1AG057892, and RF1NS136995.

## Notes

### Competing Interest Statement

The authors have declared no competing interest.

